# Lysophosphatidic acid, a CSF marker in posthemorrhagic hydrocephalus that drives CSF accumulation via TRPV4-induced hyperactivation of NKCC1

**DOI:** 10.1101/2022.01.24.477507

**Authors:** Trine Lisberg Toft-Bertelsen, Dagne Barbuskaite, Eva Kjer Heerfordt, Sara Diana Lolansen, Søren Norge Andreassen, Nina Rostgaard, Markus Harboe Olsen, Nicolas H. Norager, Tenna Capion, Martin Fredensborg Rath, Marianne Juhler, Nanna MacAulay

## Abstract

A range of neurological pathologies can cause secondary hydrocephalus. For decades, treatment has been limited to surgical CSF diversion, as still no pharmacological options exist due to the elusive molecular nature of the CSF secretion apparatus and its regulatory properties. We have now identified phospholipid lysophosphatidic acid (LPA) as a biomarker for posthemorrhagic hydrocephalus (PHH) in patients with subarachnoid hemorrhage (SAH) and mimicked our results in an animal model of intraventricular hemorrhage (IVH). Intraventricular administration of LPA caused elevated brain water content and ventriculomegaly in experimental rats, via its action as an agonist of the choroidal transient receptor potential vanilloid 4 (TRPV4) channel. TRPV4 was revealed as a novel regulator of ICP in experimental rats via its ability to modulate the CSF secretion rate through its direct activation of the Na^+^, K^+^, 2Cl^-^ cotransporter (NKCC1) implicated in CSF secretion. Together, our data reveal that a biomarker present in brain pathologies with hemorrhagic events promotes CSF hypersecretion and ensuing brain water accumulation via its direct action on TRPV4 and its downstream regulation of NKCC1. TRPV4 may therefore be a promising future pharmacological target for pathologies involving disturbed brain fluid dynamics.

## INTRODUCTION

The mammalian brain is submerged in cerebrospinal fluid (CSF), the majority of which is continuously produced by the monolayered epithelial structure, the choroid plexus, lining the fluid-filled ventricles [1, 2]. The CSF serves crucial roles such as cushioning the brain and thus protecting it from mechanical insult [3] in addition to acting as the transport route for delivery/disposal of nutrients, metabolites, and immune modulators [4]. Dysregulation of CSF dynamics occurs in a range of neuropathologies, amongst others posthemorrhagic hydrocephalus (PHH) [5–7], in which water accumulates in the brain after the hemorrhagic event, causing ventriculomegaly and elevation of the intracranial pressure (ICP). PHH may arise secondary to spontaneous or traumatic intracranial haemorrhage in all age groups [8, 9], particularly frequently following hemorrhage in the subarachnoid and ventricular compartments. Such hemorrhagic event is, in addition, often observed in the infantile hydrocephalus that occurs in approximately 1 in 1,000 newborn children [10, 11] and arises mainly due to germinal matrix bleeding of prematurity. In older children and adults, PHH is mostly due to bleeding from aneurysms or vascular malformations. In elderly patients, PHH is mostly caused by bleeding into the ventricular system from a hypertensive parenchymal haemorrhage [12–19]. Such pathological posthemorrhagic brain water accumulation is generally attributed to impaired clearance of CSF caused by obstruction of CSF flow pathways. Currently, the only available treatment relies on drainage of the excess fluid from the brain via surgical CSF diversion through external ventricular drainage, ventricular shunt placement or ventriculostomy [20, 21]. These surgical procedures associate with frequent side effects, i.e. infections and shunt malfunctions, which necessitate alternative treatment options. However, pharmacological interventions have proven suboptimal [22, 23], partly due to the knowledge gap concerning the molecular mechanisms of CSF secretion and their regulatory properties in health and disease.

Emerging evidence suggests a component of CSF hypersecretion in some forms of hydrocephalus in patients and experimental animal models, i.e. choroid plexus hyperplasia, choroid plexus tumours, or PHH [5, 6, 24]. In the rodent experimental model of intraventricular hemorrhage (IVH), the PHH was demonstrated to occur following CSF hypersecretion [6]. The molecular coupling between the hemorrhagic event and the hydrocephalus formation was proposed to rely on the associated inflammatory response promoting hyperactivity of the choroidal Na^+^/K^+^/2Cl^-^ cotransporter NKCC1 [6], that serves as a key contributor to CSF formation by rodent and canine choroid plexus [6, 25–27]. However, a component of the PHH may be contributed to the ventricular actions of the lipid, lysophosphatidic acid (LPA) [28, 29], which enters the ventricular system with the hemorrhagic event and is elevated in CSF of patients and experimental mice experiencing traumatic brain injury [30, 31].

The choroid plexus epithelial cells abundantly express the calcium-permeable nonselective cation channel transient receptor potential vanilloid 4 (TRPV4) [32, 33]. TRPV4 is polymodal in a sense of its several distinct manners of activation, one of which being lipid-mediated modulation of channel activity [34]. TRPV4 has been implicated in CSF secretion following demonstration of its ability to modulate transepithelial ion flux in choroid plexus cell lines [35, 36] and of TRPV4 antagonists effectively alleviating ventriculomegaly in a genetic animal model of hydrocephalus [37]. TRPV4 could thus act as the molecular link coupling the hemorrhage-mediated LPA elevation to the CSF hypersecretion and ensuing hydrocephalus.

Here, we demonstrate that LPA is indeed elevated in PHH patients and in an animal model thereof and reveal its ability to directly modulate the ion channel TRPV4, resulting in NKCC1-mediated CSF hypersecretion and ventriculomegaly.

## RESULTS

### LPA is elevated in PHH patients

CSF from patients with subarachnoid hemorrhage (SAH) was analysed for LPA content with a sensitive alpha-LISA assay, which allows for precise detection of solutes in small fluid quantities [38]. The LPA concentration was significantly increased in the CSF from patients with SAH (0.61 ± 0.06 ng/ml, n=13) compared to CSF obtained from control subjects during preventive surgery for unruptured aneurysms (0.44 ± 0.03 ng/ml, n=14, P<0.05, Fig. 1A), which demonstrated that SAH associates with an elevated concentration of LPA in the CSF.

**Fig.1.**
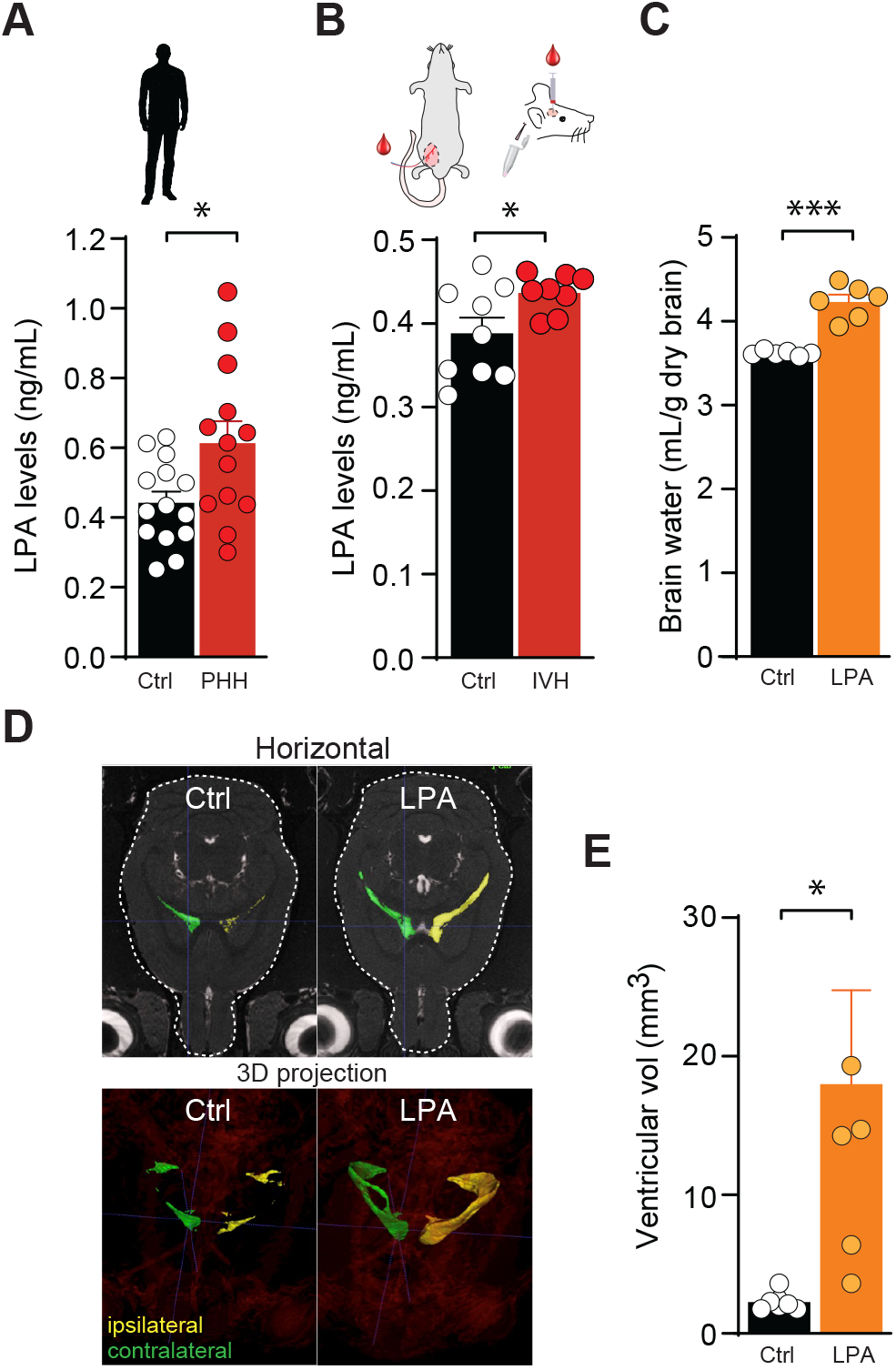
PHH associates with elevated levels of LPA, excessive brain water and ventriculomegaly. **A** Quantification of LPA levels in CSF sampled from patients with SAH, n=13 (red) and control subjects, n=14 (black). **B** Quantification of LPA levels in CSF sampled from IVH (inset) rats, n=8 (red) and sham operated rats, n=9 (black). **C** Brain water content quantified from LPA-treated rats, n=6 (orange) and sham operated rats, n=6 (black). **D** Representative MRI brain sections (upper panel, horizontal; lower panel, 3D projections) from rats 24 h post-surgery, n=3 of each. **E** Ventricular volume quantification from MRI sections from LPA-treated rats, n=3 (orange) and sham operated rats, n=3 (black) 24 h post-surgery. Statistical evaluation with Student’s t-test. * P<0.05; *** P<0.001.

### The clinical observation of elevated levels of LPA in patients with SAH is mimicked in an experimental model of intraventricular hemorrhage

To obtain an animal model of PHH with which to resolve the molecular coupling between the hemorrhagic event and hydrocephalus formation, we verified the LPA elevation observed in PHH patients in a rat model of IVH. CSF was extracted from anesthetized rats 24 h after intraventricular injection of autologous blood or sham operated and saline-injected control rats [38], Fig. 1B, and the LPA abundance was determined. The increased LPA concentration mimicked that observed in PHH patients, as it was significantly increased in the CSF from rats with intraventricular blood delivery (0.44 ± 0.01 ng/ml, n=8) compared to that obtained from sham operated rats (0.39 ± 0.02 ng/ml, n=9, P<0.05, Fig. 1B). This finding suggests that LPA elevation associates with hemorrhagic events and supports the use of an animal model of IVH to resolve the molecular mechanisms underlying PHH.

### Intraventricular LPA administration increases the brain water content and leads to ventriculomegaly

To resolve if an LPA elevation could lead to elevated brain water content, we delivered LPA directly into the lateral ventricle of anaesthetized rats and determined the brain water content 24 h after LPA administration. The total brain water content was elevated in these rats (4.23 ± 0.08 ml H_2_O/g dry brain, n=6) compared to that obtained in control rats injected with vehicle (3.62 ± 0.01 ml H_2_O/g dry brain, n=6, P<0.001, Fig. 1C). To determine the distribution of the LPA-mediated excess brain fluid, the rats underwent magnetic resonance (MR) scanning 24 h post-LPA administration, and the scans demonstrated ventriculomegaly in the rats exposed to LPA (Fig. 1D). Quantification of the ventricular volumes revealed a robust enlargement in the LPA-treated animals (18.1 ± 6.8 mm^3^, n=3) compared to the control rats (2.3 ± 0.3 mm^3^, n=3, P<0.05, Fig. 1E). These results indicate that the elevated LPA observed in hemorrhagic events triggers fluid accumulation within the ventricular system of the rat brain.

### LPA is a novel agonist of TRPV4, which is highly expressed in choroid plexus

To resolve the molecular link between LPA and brain water accumulation, we determined the ability of LPA to act as an agonist of the ion channel TRPV4, the activity of which is modulated by lipids [34] and which may be implicated in hydrocephalus formation [37]. To obtain an isolated setting in which to resolve LPA-mediated TRPV4 activation, TRPV4 was heterologously expressed in *Xenopus laevis* oocytes, which allow for overexpression of the channel of interest with low endogenous background channel activity. TRPV4 current activity was monitored with conventional two-electrode voltage clamp, which demonstrated an LPA-mediated ~3-fold increase in membrane currents in TRPV4-expressing oocytes (compare 2.2 ± 0.3 μA, n=8 in LPA-treated oocytes to 0.7 ± 0.1 μA non-treated oocytes, n=9, P<0.01 Fig. 2A-C). The LPA-induced increase in TRPV4-mediated current was abolished upon inhibition of TRPV4 with the specific TRPV4 inhibitor RN1734 (RN) [39] (0.5 ± 0.3 μA, n=8, P<0.001, Fig. 2C), demonstrating TRPV4 as the mediator of the LPA-induced membrane currents. The membrane currents of uninjected control oocytes were unaffected by LPA exposure (n =12, Fig. 2A-B). These data demonstrate that LPA acts as an agonist of TRPV4 and causes channel opening by its direct action on the channel. RNAseq analysis of rat choroid plexus revealed that TRPV4 is the only member of the vanilloid family of TRP channels expressed in this tissue at significant levels (>0.5 TPM; Fig. 2D). Among a filtered list of transcripts encoding ion channels expressed in choroid plexus, TRPV4 appeared as the 6^th^ highest-expressed ion channel (Fig. 2E). TRPV4 is localized to the luminal membrane of rat choroid plexus, as illustrated with immunohistochemistry (Fig. 2F), and verified with Western blotting of biotin-labelled choroidal luminal membrane proteins versus total choroid plexus (Fig. 2G).

**Fig.2.**
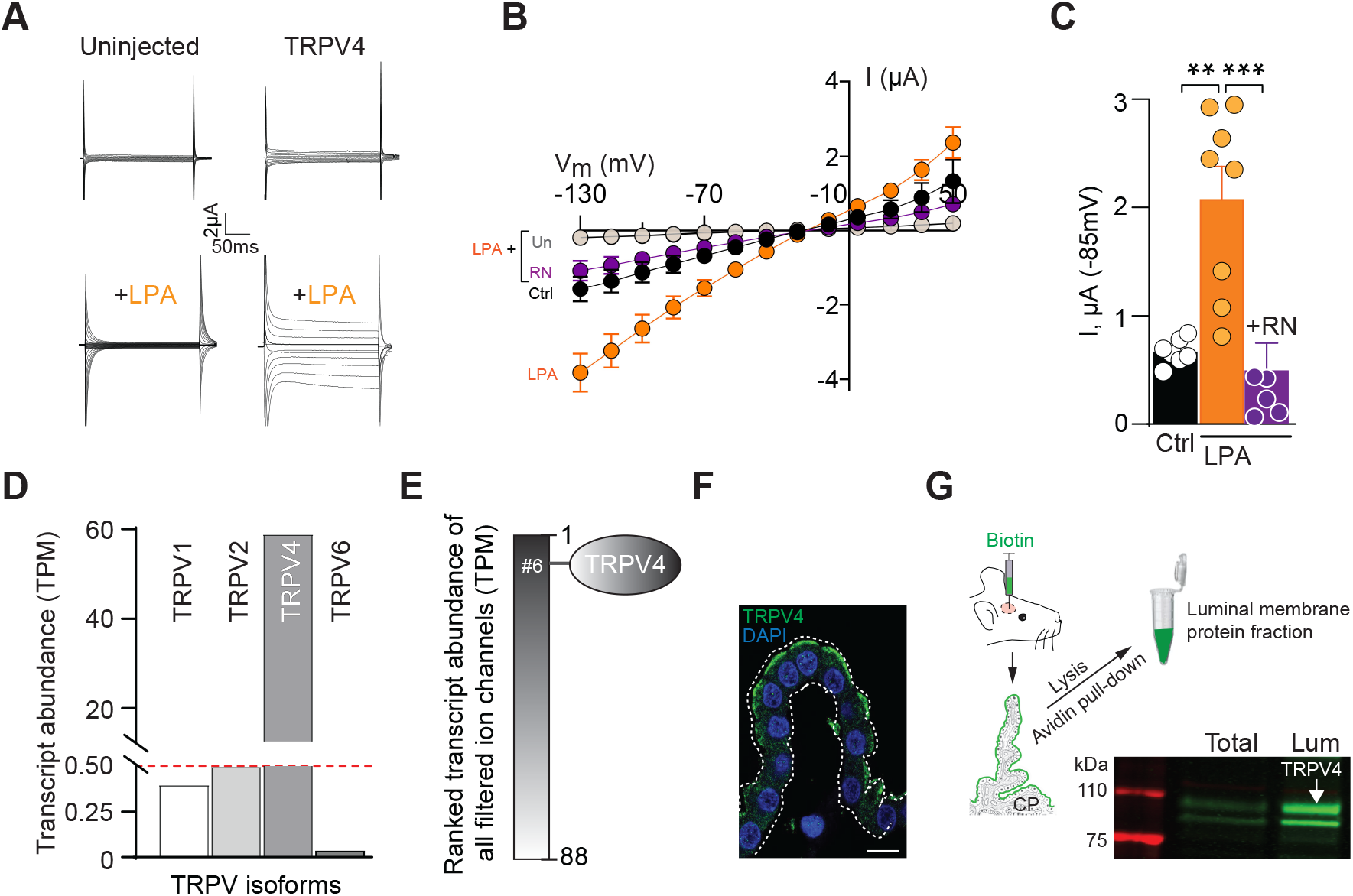
LPA is a novel endogenous agonist of luminal membraneous TRPV4. **A** Representative current traces obtained from uninjected or TRPV4-expressing oocytes during a 200 ms step protocol (upper panel, control solution; lower panel, LPA-containing solution). **B** Averaged I/V curves from TRPV4–expressing oocytes in control solution, n=9 (black), during application LPA without, n=8 (orange) or with TRPV-4 inhibition, n=8 (RN, purple). Uninjected oocytes treated with LPA are shown in grey, n=12. **C** TRPV4-mediated current activity (at V_m_ = −85 mV) summarized in control solution (black) and after exposure to LPA without (orange) or with TRPV4 inhibition (purple). **D** Transcript abundance (in TPM) of members of the transient receptor potential vanilloid family obtained from RNA sequencing of rat choroid plexus; TRPV1, 0.4; TRPV2, 0.5; TRPV4, 58.5; TRPV6, 0.06. **E** Transcript abundance of all filtered ion channels. **F** Representative immunolabeling of TRPV4 in rat choroid plexus. Nuclei staining by DAPI. Scale bar□=□5□μm. **G** Schematic illustration of biotinylation of the apical membrane of rat choroid plexus. Representative Western blot of TRPV4 expression in total tissue fraction and biotinylated membrane fraction (lum = luminal). Statistical evaluation with one-way ANOVA with Dunnett’s *post hoc* test ** P< 0.01; *** P<0.001.

### Inhibition of TRPV4 lowers the intracranial pressure

TRPV4 inhibition alleviates ventriculomegaly in a genetic model of hydrocephalus [37]. Thus, TRPV4 may well directly influence the brain water balance and modulate the ICP. We therefore monitored the ICP in anesthetized and ventilated rats during modulation of TRPV4 channel activity. ICP dynamics were recorded with a pressure probe placed in a cranial window above dura, with a continuous slow (0.5 μl/min) infusion of artificial CSF (aCSF) (containing vehicle) into one lateral ventricle (see Fig. 3A for a schematic of the experimental setup). The ICP at the start of the experiment was 4.8 ± 0.2 mmHg, n=10 (Fig. 3B inset) and reduced slightly over the course of the experiment upon mock solution change (Fig. 3B) by 7 ± 1%, n=5, Fig. 3C). Replacement of the control solution with a solution containing the TRPV4 inhibitor RN, significantly reduced the ICP (27 ± 2%, n=5) compared to that obtained in control solution (P<0.001, Fig. 3B-C). These results demonstrate that TRPV4 inhibition lowers the ICP in rats *in vivo* and thus suggest a direct involvement of the ion channel in controlling brain fluid accumulation.

**Fig.3.**
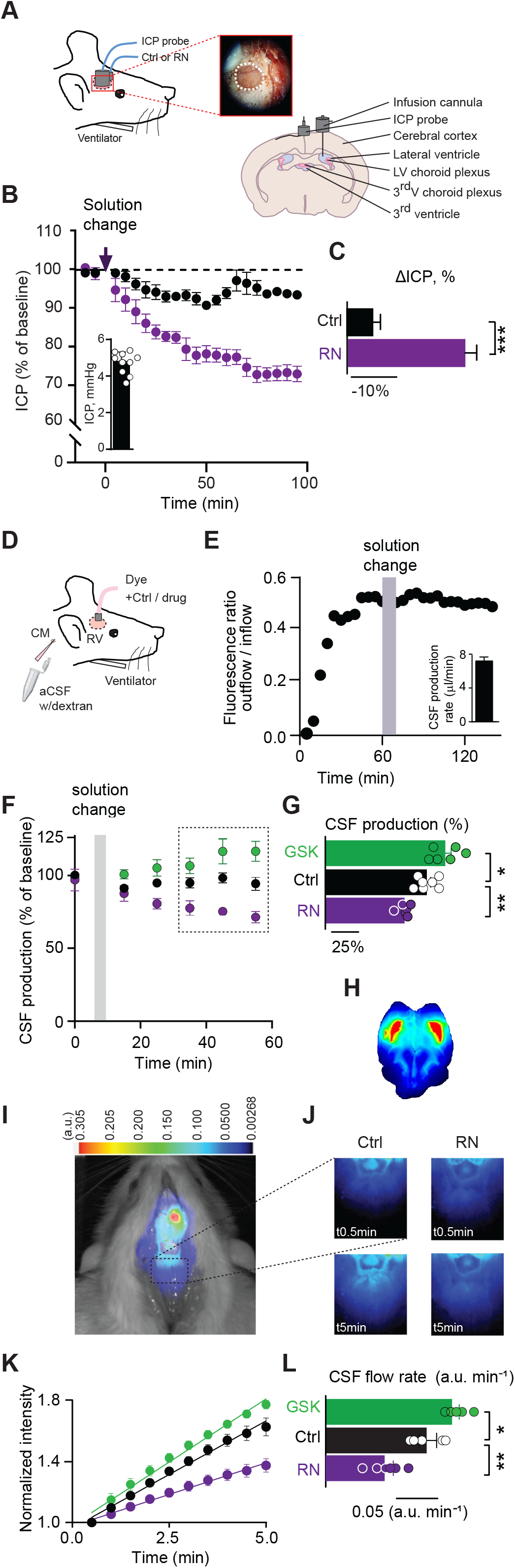
TRPV4 activity modulates ICP and CSF production *in vivo*. **A** Schematic illustrating the cranial window (shown as a dotted circle) into which the ICP probe is positioned, LV; lateral ventricle. **B** ICP as a function of time upon infusion of control solution, n=5 (black) or TRPV4 inhibitor-containing solution, n=5 (RN; purple) shown as 5 min average values normalized to the baseline. Inset, initial ICP, n=10. **C** Summarized changes in ICP with control solution (black) and TRPV4 inhibitor-containing solution after 1.5 h. **D** Schematic of the VCP method used to determine the CSF production rate. **E** Representative time course of the dextran ratio (outflow/inflow) with a mock solution change with control solution (indicated with a grey bar). Inset, average CSF production rate, n=7. **F** CSF production as a function of time. Data normalized to the last four samples before solution change to either the control solution, n=7 (black), the TRPV4 activator GSK, n=6 (green), or the TRPV4 inhibitor RN, n=6 (purple). **G** Summarized CSF production rates in % of control after exposure to vehicle (black), TRPV4 activation (green) and TRPV4 inhibition (purple). **H** Correctly targeted dye delivery in mid-saggital sections of a rat brain. **I** A representative image of a rat after injection of IRDye 800CW carboxylate dye (superimposed pseudo-color). The square placed in line with lambda indicates the area of dye content quantification. **J** Representative images obtained at t=_0.5_ (t_0.5_) and t=5 (t_5_) min in control solution (left) or upon TRPV4 inhibition (RN; right). **K** The dye intensity normalized to that obtained in the first image and plotted as a function of time representing flow rate for control, n=5 (black), TRPV4 activation, n=6 (GSK, green) and TRPV4 inhibition, n=6 (RN, purple). **L** Quantification of the dye intensity (flow rate) determined from linear regression in **K** over the 5 min. time window from control (black), TRPV4 activation (green) and TRPV4 inhibition (purple). Statistical evaluation with Student’s t-test (ICP) or one-way ANOVA with Dunnett’s *post hoc* test (VCP) * P<0.05; ** P<0.01; *** P<0.001.

### TRPV4 activity modulates the CSF production

To resolve whether the TRPV4-mediated ICP modulation originated from altered CSF production, we determined the effect of TRPV4 activity on the rate of CSF secretion in anesthetized and ventilated rats. CSF secretion was quantified by the ventriculo-cisternal perfusion (VCP) technique in which dextran-containing aCSF is perfused at a constant rate into the lateral ventricle and CSF samples collected from the cisterna magna (see Fig. 3D for a schematic). To obtain a time control experiment demonstrating the stability of the recording, the control aCSF was replaced by an identical solution after which the CSF secretion rate remained stable during the entire experimental window (95 ± 3 % of control, n=7, P=0.14, Fig. 3E-G). Dextran is cell membrane impermeable and remains within the ventricular system [27], so the CSF secretion rate is obtained by quantification of the dextran dilution that occurs upon mixing with newly secreted CSF (7.1 ± 0.7 μl/min, n = 7, Fig. 3E, inset). Inhibition of TRPV4 by inclusion of the TRPV4 inhibitor RN to the infused aCSF led to a reduction in the CSF secretion rate to 75 ± 2% of control (n=6, P<0.01) whereas activation of TRPV4 with its agonist GSK1016790A (GSK) caused an elevation of the CSF secretion (to 113 ± 6 % of control, n=6, P<0.05), Fig. 3F-G. These data illustrate that modulation of the TRPV4 activity affects the rate of CSF secretion in anesthetized rats. With the long experimental duration of the VCP experiments, extrachoroidal TRPV4 could be affected and indirectly modulate CSF secretion e.g. via change in cardiovascular parameters and nerve activity. To obtain a swifter, and less invasive, protocol for determining the rate of CSF secretion, we employed whole animal live imaging of CSF dynamics based on a fluorescent tracer. A fluorescent dye (IRDye 800CW) was injected into the lateral ventricle of an anesthetized rat (see example of correctly targeted dye delivery in Fig. 3H), which was immediately placed in the fluorescent scanner. Recording of the caudal redistribution of the fluorescent dye initiated within 1 min of dye injection served as a proxy of CSF production. The dye movement is seen as superimposed pseudo-color fluorescence (Fig. 3I-J) and the dye intensity quantified in a region of interest placed in line with lambda (Fig. 3I). Intraventricular delivery of the TRPV4 inhibitor RN reduced the dye movement by 38% (from 0.13 ± 0.01 a.u. min^-1^, n=5 in control animals to 0.08 ± 0.01 a.u. min^-1^, n=6, P<0.01), while inclusion of the TRPV4 agonist GSK increased the flow rate 23% (to 0.16 ± 0.01 a.u. min^-1^, n=6, P<0.05), Fig. 3K-L. Taken together, these findings reveal that TRPV4 activity directly modulates the CSF secretion rate.

### TRPV4 co-localizes with NKCC1 and modulates the CSF secretion apparatus by NKCC1 activation

With a polarization to the luminal membrane, TRPV4 could exert its stimulatory effect on CSF secretion by modulation of the cotransporter NKCC1 that is implicated in CSF secretion [6, 25, 26]. Co-localization of TRPV4 and NKCC1 in the luminal membrane of choroid plexus was demonstrated by immunohistochemistry (Fig. 4A). Their close proximity (30-40 nm) was verified with the proximity ligation assay [40], in which fluorescent speckles occur solely when the respective antibodies are in close proximity, as demonstrated in (Fig. 4B, left panel). Fluorescent speckles were absent if the primary antibodies was omitted (Fig. 4B, right panel). These results indicate that TRPV4 and NKCC1 are closely co-localized on the luminal membrane of the choroid plexus. To reveal a functional coupling between TRPV4 and NKCC1, we quantified the NKCC1 transport activity with *ex vivo* radioisotope efflux experiments in acutely isolated rat choroid plexus pre-loaded with ^86^Rb^+^, which serves as a congener for K^+^ (Fig. 4C inset). TRPV4 activation by GSK increased the rate of ^86^Rb^+^ efflux ~4 fold (from 0.35 ± 0.06, n=5, in control to 1.32 ± 0.15, n=5, P<0.001, Fig. 4C). A direct effect on NKCC1-mediated efflux was ascertained with inclusion of the NKCC1 inhibitor bumetanide [41], which reduced the GSK-mediated efflux (to 0.71 ± 0.09, n=5, Fig. 4C). These data illustrated a GSK-mediated ~3 fold increase in the bumetanide-sensitive part of the ^86^Rb^+^ efflux (0.22 ± 0.05 min^-1^ compared to 0.61 ± 0.06 min^-1^, Fig. 4D, P<0.001), which is thus assigned to elevated NKCC1 activity. Verification of the TRPV4-mediated increase in the NKCC1-mediated efflux rate was obtained with attenuation of the GSK-mediated elevation of the ^86^Rb^+^ efflux by inclusion of the TRPV4 inhibitor, RN (to 0.66 ± 0.02 min^-1^, n=5). These data demonstrate a robust TRPV4-mediated increase in NKCC1 activity in choroid plexus.

**Fig.4.**
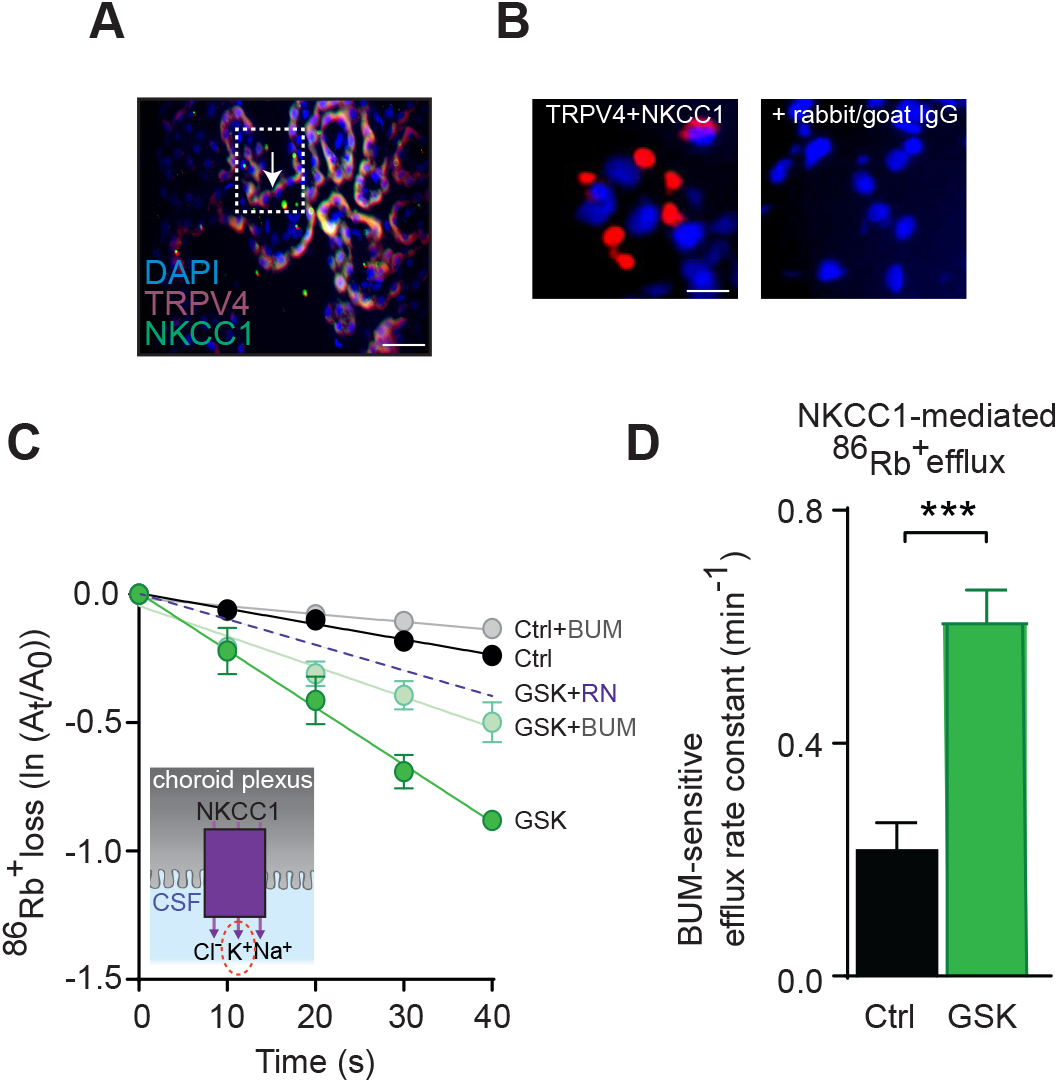
TRPV4 co-localizes with – and regulates – NKCC1. **A** Representative immunolabeling in rat choroid plexus of TRPV4 (red) and NKCC1 (green). DAPI is used for nuclei staining. Scale bar□ = 25 ≥ μm. **B** Proximity ligation assayed interaction complex between NKCC1 and TRPV4 shown as red speckles (left). Proximity ligation in the absence of primary antibodies against TRPV4 and NKCC1. Scale bar = 10 μm. **C** Efflux of ^86^Rb^+^ from choroid plexus (inset) in control settings without, n=5 (black) or with NKCC1 inhibition by bumetanide (BUM), n=5 (grey), or upon TRPV4 activation by GSK without, n=5 (green) or with BUM, n=5 (light green). GSK-mediated ^86^Rb^+^ efflux obtained with inclusion of the TRPV4 inhibitor RN, n=5 (dotted, purple line). Y-axis is the natural logarithm of the amount of left in the choroid plexus at time t (A_t_) divided by the amount at time 0 (A_0_). **D** NKCC1-mediated efflux rate constants for ^86^Rb^+^ in control, n=5 (black) or upon TRPV4 activation, n=5 (GSK; green). Statistical evaluation with Student’s t-test.*** P<0.001.

### TRPV4-mediated activation of NKCC1 in choroid plexus requires intracellular Ca^2+^ dynamics

To identify the molecular pathway coupling TRPV4 channel activity to NKCC1 activation, we monitored the TRPV4-induced Ca^2+^ dynamics in acutely isolated choroid plexus *ex vivo.* The tissue was loaded with the Ca^2+^-sensitive fluorescent dye Fluo-8 AM, which allowed for recording of the spontaneous global Ca^2+^ dynamics (Fig. 5A). Exposure to the TRPV4 activator GSK caused a global increase in intracellular [Ca^2+^] (from 1272 ± 170 a.u. to 1907 ± 134 a.u., n=5, P<0.05, Fig. 5B-C). To reveal whether these TRPV4-mediated Ca^2+^ fluctuations are permissive for the NKCC1 activation, we determined the NKCC1 activity, assessed by ^86^Rb^+^ efflux experiments on *ex vivo* rat choroid plexus, in the absence of Ca^2+^ (Ca^2+^-free test solutions and chelation of [Ca^2+^]_i_ with BAPTA-AM). The ^86^Rb^+^ efflux in the presence of GSK (1.41 ± 0.12 min^-1^, n=4) was reduced in the absence of Ca^2+^ (0.78 ± 0.07 min^-1^, n=4, P<0.01), Fig. 5D-E. The GSK-mediated ^86^Rb^+^ efflux was thus halved in the absence of Ca^2+^ (1.10 ± 0.09 min^-1^ compared to 0.53 ± 0.05 min^-1^, Fig. 5E), suggesting that NKCC1 is activated by TRPV4 via its ability to cause Ca^2+^ accumulation in the choroid plexus upon channel activation.

**Fig.5.**
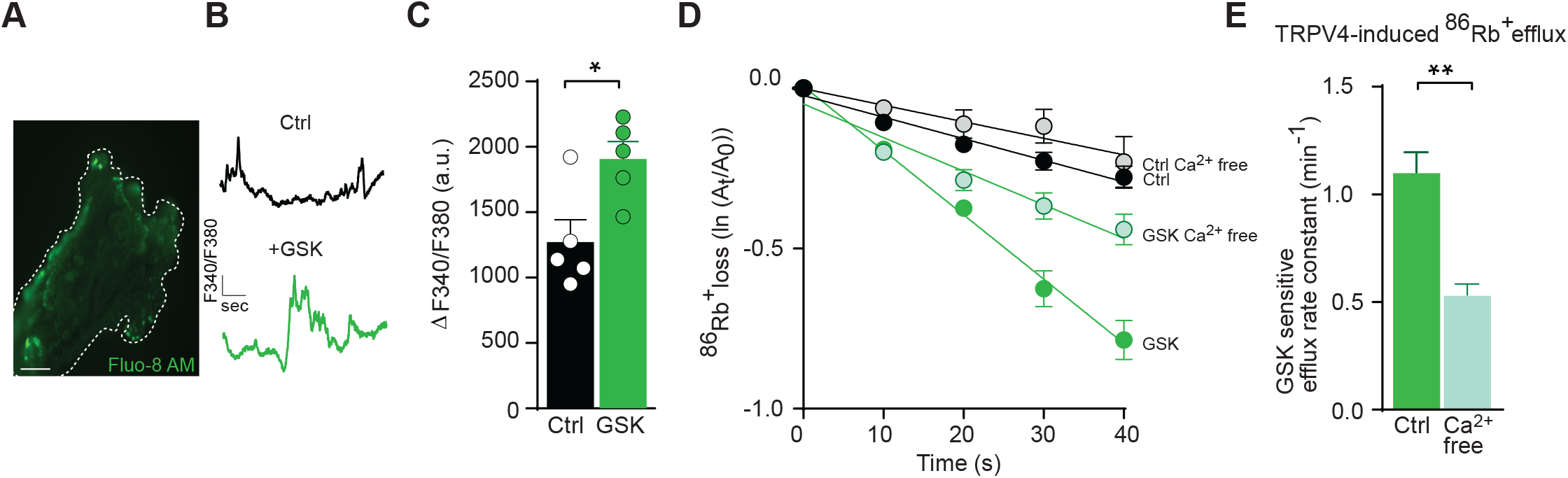
Activation of TRPV4 in choroid plexus epithelial cells induces altered Ca^2+^ dynamics. **A** Micrograph of a rat choroid plexus loaded with Fura-2AM. The dotted line indicates the outline of the tissue. Scale bar□= 400 μm. **B** Representative raw traces illustrating [Ca^2+^]_i_ dynamics in choroid plexus (control, black; GSK-induced activation of TRPV4, green). **C** Averaged Fura-2AM intensity in choroid plexus exposed to control solution, n=5 (black) or GSK-containing solution, n=5 (green). **D** Efflux of ^86^Rb^+^ from choroid plexus in control solution, n=4 (black), in Ca^2+^-free solution, n=4 (grey), upon TRPV4-activation by GSK with, n=4 (green) or without Ca^2+^, n=4 (light green). **E** GSK-sensitive efflux rate constants for ^86^Rb^+^ with (green) or without (light green) Ca^2+^ present. Statistical evaluation with Student’s t-test. *P<0.05; **P<0.01.

### TRPV4 activates NKCC1 in a WNK-SPAK-dependent manner

Ca^2+^ fluctuations may regulate the WNK and SPAK kinases [42], the latter of which has been implicated in NKCC1-mediated hypersecretion of CSF (Fig. 6A) [6]. RNA-Seq from rat choroid plexus revealed SPAK as the most abundantly expressed kinase, with WNK1 placing at rank 74, WNK2 at rank 130 and WNK4 at rank 240 out of the 297 kinases detected in the RNA-seq data (Fig. 6B). Network analysis illustrates established links between NKCC1 and the kinases with a single putative connection to TRPV4 (Fig. 6C). The TRPV4/NKCC1/WNK1/4 have a high confident score (>0.7 string score), whereas TRPV4 is only connected to the network through co-mentions in the literature with medium string score of 0.51. To determine whether TRPV4-mediated Ca^2+^ dynamics activate NKCC1 in a WNK and/or SPAK-dependent fashion, we determined GSK-mediated choroidal NKCC1 activity, by *ex vivo* ^86^Rb^+^ efflux experiments, in the presence of inhibitors of these kinases. Inhibition of SPAK and WNK did not affect the basal NKCC1 activity (Fig. 6D+F), but the SPAK inhibitor closantel [6], strongly reduced the GSK-mediated ^86^Rb^+^ efflux (from 0.81 ± 0.04 min^-1^, n=6 to 0.15 ± 0.03 min^-1^, n=6, P<0.001), Fig. 6D-E. The pan-WNK inhibitor WNK463 [43] reduced the GSK-mediated ^86^Rb^+^ efflux (from 0.87 ± 0.16 min^-1^, n=5 to 0.39 ± 0.07 min^-1^, n=5, p<0.01), Fig. 6F-G. The GSK-mediated ^86^Rb^+^ efflux was thus more than halved by WNK inhibition and completely abolished by SPAK inhibition. These data demonstrate that NKCC1 is activated by TRPV4 via Ca^2+^-mediated activation of the WNK/SPAK signalling pathway.

**Fig.6.**
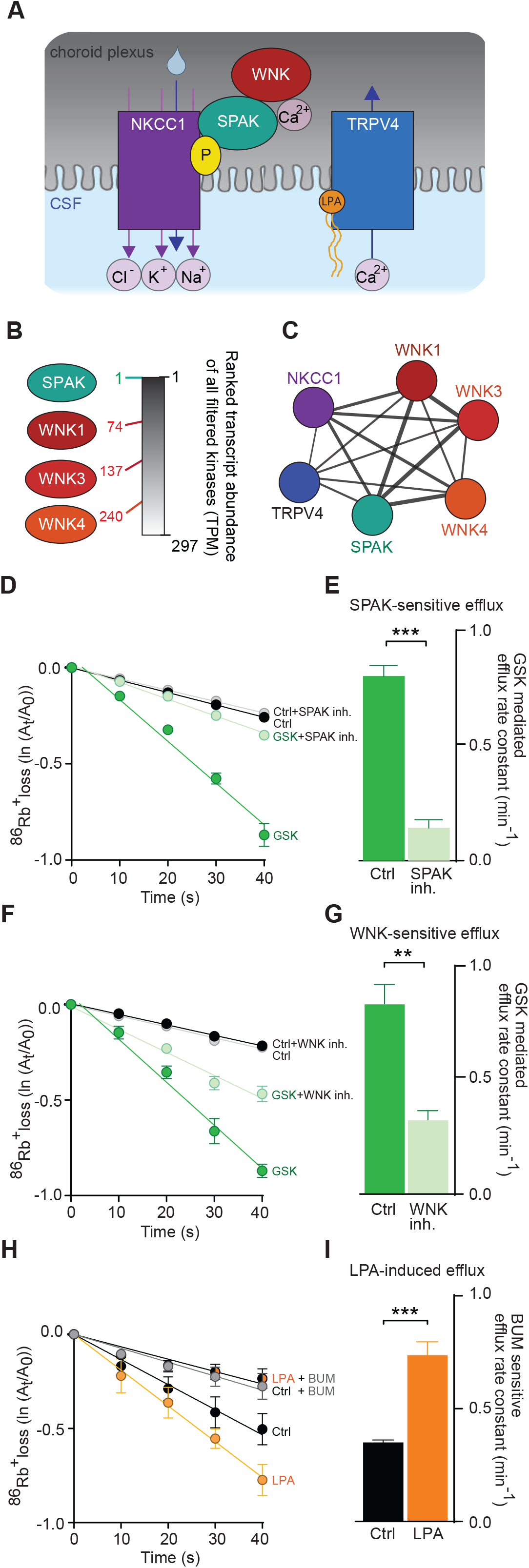
LPA-induced activation of TRPV4 regulates NKCC1-mediated CSF efflux in a WNK-SPAK-dependent fashion. **A** Schematic of the hypothesized LPA-induced TRPV4-mediated Ca^2+^ influx regulating WNK-SPAK-mediated phosphorylation of NKKC1. **B** Transcript abundance of filtered kinases in rat choroid plexus with rank of SPAK and WNK kinases depicted. **C** Network analysis depicting published protein-protein associations amongst those illustrated in Panel A. **D** ^86^Rb^+^ efflux from choroid plexus in control settings without, n=6 (black) or with SPAK inhibition, n=6 (grey), or upon TRPV4 activation by GSK without, n=6 (green) or with, n=6 (light green) SPAK inhibition. Y-axis is the natural logarithm of the amount of left in the choroid plexus at time t (A_t_) divided by the amount at time 0 (A_0_). **E** GSK-mediated efflux rate constants for ^86^Rb^+^ without (green) or with SPAK-inhibition (light green). **F** ^86^Rb^+^ efflux in control settings without, n=5 (black) or with, n=5 (grey) WNK inhibition, or upon TRPV4 activation by GSK without, n=5 (green) or with, n=5 (light green) WNK inhibition. **G** GSK-mediated efflux rate constants for ^86^Rb^+^ in without (green) or with (light green) WNK-inhibition. **H** ^86^Rb^+^ efflux in control settings without, n=5 (black) or with, n=5 (grey) NKCC1 inhibition by bumetanide (BUM), or upon application of LPA without, n=5 (orange) or with, n=5 (black/orange) NKCC1 inhibition (BUM). **I** BUM-sensitive efflux rate constants for ^86^Rb^+^ in control setting (black) or upon LPA application (orange). Statistical evaluation with Student’s t-test.** P<0.01; *** P<0.001.

### LPA activates NKCC1 in choroid plexus

With the demonstration of LPA-mediated TRPV4 activation (Fig. 2) and TRPV4-induced NKCC1 activation (Fig. 4) promoting CSF hypersecretion (Fig. 4), we verified LPA-mediated hyperactivity of NKCC1 in choroid plexus by assessing the ^86^Rb^+^ efflux rate in the presence of this lipid. LPA elevated the efflux rate (from 0.36 ± 0.01 min^-1^, n=5 to 0.74 ± 0.05 min^-1^, n=5, P<0.001, Fig. 6H) in a bumetanide-sensitive manner, which demonstrated a ~2-fold increase in NKCC1 activity upon inclusion of LPA (Fig. 6I). LPA is thus indeed able to act *in situ* and mediate NKCC1 hyperactivity in choroid plexus.

## DISCUSSION

Here, we reveal a molecular coupling between a brain hemorrhagic event and the ensuing ventricular enlargement signifying PHH. The origin of hydrocephalus formation is generally sought in blockage of the CSF exit routes [44–47]. In conditions with no discernable exit route blockage, ventriculomegaly could be caused by CSF hypersecretion [6, 48–50]. Hypersecretion of CSF may occur particularly in hydrocephalus etiologies where inflammation could be a pathogenetic factor [6, 48, 51]. However, here we report that the mere presence of a blood lipid, LPA, appears to cause CSF hypersecretion leading to ventriculomegaly in experimental rats. We detected an elevation of the phospholipid LPA in the CSF from patients with SAH and from rats following experimentally-induced IVH. Such LPA elevation has previously been reported for patients and mice experiencing traumatic brain injury [30, 31], indicative of brain entrance of the serum lipid LPA along with the hemorrhagic event. Mouse models of neonatal (embryonic or postnatal) PHH displayed a range of LPA-mediated morphological changes in the week(s) following the intrauterine intraventricular LPA administration, such as ventricular enlargements, 3^rd^ ventricular occlusion, elevated ICP, thinning of cortical layers, and cilia loss along the lateral ventricular walls [28, 29]. LPA effectuated these features via their G protein-coupled LPA receptors expressed in the brain tissue (LPA_R1_-LPA_R6_) [28, 29, 52]. Here, we demonstrate an additional acute (24 hour) effect of intraventricularly-delivered LPA causing ventriculomegaly and elevated brain fluid content in adult rats by its ability to act as an agonist of the TRPV4 channel. TRPV4 is expressed in various tissues and cell types throughout the body, for review see [34], but in the brain it is highly concentrated in the choroid plexus [35, 37, 53], in which we reveal it as the 6^th^ highest-expressed ion channel, localized to the luminal membrane (this study and [33]). Inhibition of TRPV4 modulates transepithelial ion flux in immortalized choroid plexus cell lines [35, 36] and abolishes ventriculomegaly in a genetic rat model of hydrocephalus [37]. Accordingly, we demonstrated an acute reduction of ICP in healthy rats exposed to an intraventricularly-delivered TRPV4 inhibitor. The reduction in ICP came about by TRPV4’s ability to directly modulate the rate of CSF secretion. TRPV4 activity thus modulates the CSF secretion rate in healthy rats and thereby governs the ICP. Notably, with its luminal localization on the choroid plexus, TRPV4 inhibition may fail to reach its target if non-cell permeable inhibitors are delivered intraperitoneally (i.p.) [54].

TRPV4 can be directly activated by various lipid compounds [34], and we here add LPA to the list of lipid agonists of TRPV4. LPA exposure elevated the TRPV4-mediated current in TRPV4-expressing *Xenopus laevis* oocytes, an elevation that was absent in the presence of a TRPV4 inhibitor and in control oocytes lacking TRPV4 expression. Such a reduced experimental system allows for determination of direct functional interactions and promoted LPA as a direct agonist of TRPV4. The molecular coupling between TRPV4 activation and elevated CSF secretion originated in the TRPV4-mediated Ca^2+^ influx, which activated the WNK/SPAK signaling pathway as also observed in salivary glands [42]. These kinases are highly expressed in choroid plexus, with SPAK ranking as the highest expressed kinase (this study), and are well-established regulators of NKCC1 activity via modulation of the transporter phosphorylation status [6, 55, 56]. Activation of this pathway, with either LPA or a well-established synthetic lipid agonist of TRPV4, terminated in NKCC1 hyperactivity and a resultant elevation of the CSF secretion rate (see schematic in Fig. 6). NKCC1 is a key contributor to the CSF secretion in mice, rats, and dogs [6, 25, 26] as revealed with intraventricular delivery of the NKCC1 inhibitor bumetanide. With alternative delivery routes (i.v. or i.p.), bumetanide fails to reach its target on the luminal surface of choroid plexus in sufficient concentrations [6, 54]. A coupling between the SPAK/WNK signaling cascade and NKCC1-mediated CSF hypersecretion has previously been described in an experimental IVH rat model [6]. That study, however, promoted the toll-like receptor 4 and NFκβ as the molecular links between the hemorrhagic event and the SPAK-induced NKCC1-mediated CSF hypersecretion [6]. These two PHH-related molecular pathways both promoting NKCC1 hyperactivity could well occur in parallel on different time scales with the LPA/TRPV4-mediated effects discernible within minutes, followed by a slower route (minutes-hours) through a hemorrhage/TLR4-mediated path through NFκβ-mediated transcription events leading to activation of SPAK [6]. On a more prolonged time scale (days-weeks) LPA may act on its various receptors and promote the ciliopathy, 3^rd^ ventricular occlusion, and potentially other micro-blockages not discernible on brain imaging. PHH may thus occur by various mechanisms and in distinct structures in the brain at different time points following the hemorrhagic event [48]. Importantly, it appears that PHH arises, in part, from a component of CSF hypersecretion (this study and [6]), which may well occur in other brain pathologies with disturbed brain fluid dynamics. Delineation of the molecular underpinnings governing this (and other) CSF secretory disturbances may open new avenues for the pharmacological therapy wanting for these conditions. Although more could arise with future studies, potential targets could be TRPV4 [37], the LPA signalling pathways [30], the NKCC1 [6], or its regulatory kinase, SPAK [57, 58], inhibition of which may reduce hydrocephalus or lesion size in different cerebral pathological insults.

Limitations to the study include CSF sampling from cisterna magna in the rodent experimentation, from the ventricular compartment in the patients with SAH, and from the basal cisterns in the control patients. The different patient sample sites were dictated by ethical limitations in invasive CSF sampling, but could influence our results if the CSF composition differs between these locations. The control samples may contain blood contamination from the surgical opening. Such contamination would raise the LPA levels in the control CSF. Our results would thus be expected to show even further differences between the two CSF groups, if one could test control CSF samples with no traces of blood. Of note, with the invasive nature of experimental approaches towards determination of agonist-induced modulation of intracranial pressure, cerebrospinal fluid secretion rates, and choroidal transport mechanisms, these parameters were obtained from the rodent animal model. Such findings may not fully capture the human condition.

In conclusion, we demonstrate that the serum lipid LPA, entering the ventricular system during a hemorrhagic event, leads to CSF hypersecretion and the ensuing ventriculomegaly signifying PHH. LPA acts directly on TRPV4, as a novel lipid agonist of this choroidal ion channel, the activation of which causes the NKCC1 hyperactivity underlying the TRPV4-mediated elevated CSF secretion rate. Future studies aimed at elucidating the pathophysiological changes in brain fluid dynamics occurring with diverse neuropathologies, ideally, should take into account formation rates of CSF, its path through the ventricular system, as well as its drainage, as most of these pathologies are likely to affect more than one of these aspects.

## Acknowledgements

We thank laboratory manager Trine Lind Devantier and technician Rikke Lundorff, the Core Facility for Integrated Microscopy and the Panum NMR Core Facility, Faculty of Health and Medical Sciences, University of Copenhagen.

## Author contributions

Conception and design of research: T.L.T.B. and N.M.; Patient contact and CSF sampling: M.J., N.R., M.H.O., N.H.N., T.B.C.; Conduction of the experiments: T.L.T.B., D.B., E.K.H., S.D.L., S.N.A., M.F.R.; Analysis of data: T.L.T.B., D.B., E.K.H., S.D.L., S.N.A.; Interpretation of results: T.L.T.B. and N.M.; Preparation of figures: T.L.T.B.; Drafting of manuscript: T.L.T.B. and N.M.; Revision and approval of manuscript: T.L.T.B., D.B., E.K.H., S.D.L., S.N.A., N.R., M.H.O., N.H.N., T.C., M.F.R., M.J., and N.M.

## METHODS

### Patients

CSF samples were collected between June 20Í9 and September 2021 from 13 patients (mean age: 63y, range: 40-77 y, 8 F/ 5 M) with acute SAH admitted and treated for the condition at Department of Neurosurgery at Rigshospitalet, Copenhagen, Denmark. CSF samples were obtained within 24 h of ictus (n=8) or as soon as possible hereafter (n=5) through an external ventricular drain (EVD) inserted on clinical indications. To exclude that measured CSF parameters could be affected by neuroinfections requiring antibiotic treatment, patients with no signs of neuroinfection at admission or during their treatment were selected. All included patients later received a permanent ventriculo-peritoneal shunt because of continued need for CSF diversion. As control subjects, 14 patients undergoing preventive surgery for unruptured aneurysms (vascular clipping) were enrolled (mean age: 61y, range: 39-71y, 8 F/5 M), and CSF was collected from the basal cisterns during surgery prior to clipping of the aneurysm. Written informed consent were obtained from all patients or next of kin depending on the capacity of the patients and the study was approved by the Ethics committee of the Capital Region of Denmark (H-1900174/69197/H-17011472).

### Animals

All animal experiments complied with the relevant ethical regulations and conformed to European guidelines. The Danish Animal Experiments Inspectorate approved the experiments (permission no. 2016-15-0201-00944 and 2018-15-0201-01595). Adult male Sprague Dawley rats (Janvier Labs) of 9 weeks used for the animal experimentation were housed with a 12:12 light cycle and access to water and food *ad libitum* accordingly to the ARRIVE guidelines.

### Anaesthesia and physiological parameters

The experimental animals were anaesthetized with intraperitoneal (i.p.) injection with 6 mg ml^-1^ xylazine + 60 mg ml^-1^ ketamine (ScanVet) in sterile water (0.17 ml/100 g body weight, pre-heated to 37°C). Animals were re-dosed with half ketamine dose as required to sustain anesthesia. Isofluorane 1000 mg/g, ScanVet) was employed for survival procedures (mixed with 1.8 1 min^-1^ air/0.1 1 min^-1^ O_2_ (Attene vet), 5% to induce anesthesia and 1-2.5% to sustain anesthesia and scanning of the animals (anesthesia was maintained at b 1-5% isoflurane in a 1/1 mixture of air/oxygen). The body temperature of the anesthetized rats was maintained at 37°C by a homeothermic monitoring system (Harvard Apparatus). Mechanical ventilation was included for anesthetic protocols longer than 30 min to ensure stable respiratory partial pressure of carbon dioxide and arterial oxygen saturation and thus stable plasma pH and electrolyte content. A surgical tracheotomy was performed and the ventilation controlled by the VentElite system (Harvard Apparatus) by 0.9 l min^-1^ humidified air mixed with 0.1 l min^-1^ O_2_ adjusted with approximately 3 ml per breath, 80 breath min^-1^, a Positive End-Expiratory Pressure (PEEP) at 2 cm, and 10% sigh for a ~400 g rat. The ventilation settings were optimized for each animal using a capnograph (Type 340, Harvard Apparatus) and a pulse oximeter (MouseOx® Plus, Starr Life Sciences) after system calibration with respiratory pCO_2_ (4.5 - 5 kPa), pO_2_ (13.3 - 17.3 kPa), and arterial oxygen saturation (98.8 - 99.4%) (ABL90, Radiometer). For survival procedures, the rats were preoperatively given the analgesics buprenorphine p.o. (0.4 mg/kg^-1^, Sandoz) and carprofen subcutaneously (5 mg/kg, Norbrook). The former was re-administered 24 h postoperatively.

### Solutions and chemicals

The majority of the experiments were conducted in HCO_3_^-^-containing artificial cerebrospinal fluid (aCSF; (in mM) 120 NaCl, 2.5 KCl, 2.5 CaCl_2_, 1.3 MgSO_4_, 1 NaH_2_PO_4_, 10 glucose, 25 NaHCO_3_, pH adjusted with 95% O_2_/5% CO_2_). In experiments where the solution could not be equilibrated with 95% O_2_/5% CO_2_, during the experimental procedure (intracranial pressure monitoring, tissue lysis for Western blotting and RNA sequencing), the solution was instead buffered by HEPES (HEPES-aCSF; (in mM) 120 NaCl, 2.5 KCl, 2.5 CaCl_2_, 1.3 MgSO_4_, 1 NaH_2_PO_4_, 10 glucose, 17 Na-HEPES, adjusted to pH 7.4 with NaOH). Pharmacological inhibitors were dissolved in DMSO and kept as stock solutions at −20°C. These were either purchased from Sigma (bumetanide: B3023, GSK1016790A: G0798, RN-1734: R0658, lysophosphatidic acid: L7260, Closantel: 34093) or MedChemExpress (WNK463: HY-100828). All control solutions contained appropriate concentrations of vehicle (DMSO, D8418, Sigma).

### Experimental intraventricular hemorrhage (IVH) in rats

The surgery was performed on rats anesthetized with isoflurane (5-2%) under aseptic conditions with body temperature maintained at 37°C using a rectal probe and feedback-controlled heating pad (Harvard Apparatus). Rats were positioned in a stereotaxic frame (Harvard Apparatus) and the skull exposed with a midline incision. A cranial burr hole was drilled above the right lateral ventricle (0.6 mm posterior and 1.6 mm lateral to bregma), after which the rats were removed from the stereotaxic frame, the femoral artery catheterized, and approximately 300 μl blood was collected (the control rats underwent sham operation). Immediately thereafter, 200 μl of this autologous blood sample (or saline) was manually injected over the course of 15 min via a 27-gauge needle inserted stereotaxically into the burr hole in the right lateral ventricle (4.5 mm ventral). The needle was kept in place for 5 min before retraction to prevent backflow. The skin incisions were closed with sutures and the rats were allowed to recover before returning to the housing facility.

### CSF extraction and Alpha-LISA

24 h post-IVH surgery, the rats were anesthetized and placed in a stereotaxic frame. CSF was sampled from cisterna magna with a glass capillary (30-0067, Harvard Apparatus pulled by a Brown Micropipette puller, Model P-97, Sutter Instruments) placed at a 5° angle (7.5 mm distal to the occipital bone and 1.5 mm lateral to the muscle-midline). CSF was collected in polypropylene tubes (Sarstedt), centrifuged at 2000 x *g* for 10 min at 4°C, divided into aliquots, and stored at −80°C. The LPA content of the CSF samples was determined with Alpha-LISA by use of a ready-to-use microwell kit designed to detect native LPA in either rats (MBS774994, MyBioSource) or humans (MBS707296, MyBioSource). The CSF samples were added to wells pre-coated with LPA antibody, followed by addition of streptavidin-horseradish peroxidase to form an immune complex. Upon a wash step, chromogen substrate solutions were added and plate reading conducted in a microplate photometer (Synery^TM^ Neo2 Multi-mode Microplate Reader; BioTek Instruments) according to the manufacturer’s instructions.

### Determination of brain water content following intraventricular LPA exposure

25 μl LPA (100 μM) was delivered intraventricularly into anesthetized rats (sham operated rats were used as comparison) as described for the blood delivery in the IVH procedure. 24 h post-injection, the rat was sacrificed and its brain swiftly removed, placed in a pre-weighed porcelain evaporating beaker (Witeg) and weighed within min after brain isolation. The brain tissue was homogenized with a steel pestle and dried at 100 °C for 72 h to a constant mass. The dry brain was weighed, and the brain water content determined in ml/gram dry weight using the equation: (wet weight - dry weight)/dry weight. The weighing was done in a randomized and blinded fashion.

### Magnetic resonance imaging (MRI)

Anesthetized rats underwent MRI in a 9.4 Tesla preclinical horizontal bore scanner (BioSpec 94/30 USR, Bruker BioSpin) equipped with a 240 mT/m gradient coil (BGA-12S, Bruker) at the Preclinical MRI Core Facility, University of Copenhagen. The scanner was interfaced to a Bruker Avance III console and controlled by Paravision 6.1 software (Bruker). Imaging was performed with an 86 mm-inner-diameter volume resonator and a 4-channel surface quadrature array receiver coil. The animal body temperature was maintained at 37 ± 0.5°C with a thermostatically controlled waterbed and its respiratory rate monitored by an MR-compatible monitoring system (SA Instruments). The imaging protocol consisted of T2-weighted 2D rapid acquisition with relaxation enhancement (2D-RARE) for reference spatial planning with the following settings: repetition time (TR) = 4000 ms, effective echo time (TE) = 60 ms, number of averaging (NA) = 4, RareFactor = 4, slice thickness = 500 μm, in-plane resolution = 137 x 273 μm, 25 coronal slices, total acquisition time (TA) = 8.5 min. For obtaining high resolution CSF volumetry, a 3D constructive interference steady-state sequence (3D-CISS) [59] image was calculated as a maximum intensity projection (MIP) from 4 realigned 3D-TrueFISP volumes with 4 orthogonal phase encoding directions (TR = 4.6 ms, TE = 2.3 ms, NA = 1, Repetitions = 2, Flip angle = 50°, 3D spatial resolution 100 x 100 x 100 μm, RF phase advance 0, 180, 90, 270°, TA = 28 min). To obtain optimal spatial uniformity, all acquired 3D-TrueFISP volumes were motion-corrected before calculation as MIP, and the image bias field was removed with Advanced Normalization Tools (ANTs) [60, 61]. For each brain sample, the total brain volume was automatically segmented by using region growing with ITK-snap (version 3.8.0) [62]. In addition, the pixel intensity factorized semi-automatic thresholding was performed to segment the lateral ventricle in each hemisphere. The volume measurement of the whole brain and lateral ventricles were performed in ITK-snap. The analysis was carried out in a blinded fashion.

### ICP monitoring

A burr hole was drilled in the right lateral ventricle (using the coordinates 1.3 mm. posterior to Bregma, 1.8 mm. lateral to the midline, and 0.6 mm ventral through the skull) of anesthetized and ventilated rats placed in a stereotaxic frame, in which a 4 mm brain infusion cannula (Brain infusion kit 2, Alzet) was placed. On the contralateral side of the skull, an epidural probe (PlasticsOne, C313G) was placed in a cranial window (4 mm diameter) and secured with dental resin cement (Panavia SA Cement, Kuraray Noritake Dental Inc.). The cannula was pre-filled with HEPES-aCSF and connected to a pressure transducer (APT300) and a transducer amplifier module TAM-A (Hugo Sachs Elektronik). The pressure signal was visualized and recorded with a sample rate of 1 kHz using BDAS Basic Data Acquisition Software (Harvard Apparatus, Hugo Sachs Elektronik). Jugular compression was applied in the beginning and at the end of the experiment to confirm proper ICP recording. Pre-heated aCSF (containing DMSO or 1.8 mM RN1734; expected ventricular concentration 50 μM) was slowly infused (0.5 μl min^-1^) into the lateral ventricle.

### CSF production rate

The CSF production rate was determined with the ventricular-cisternal perfusion (VCP) technique. An infusion cannula (Brain infusion kit 2, Alzet) was stereotaxically placed in the right lateral ventricle of an anesthetized and ventilated rat (using the same coordinates as for ICP), through which a pre-heated (37°C, SF-28, Warner Instruments) dextran-containing solution (HCO_3_^-^-aCSF containing 1 mg/ml^-1^ TRITC-dextran (tetramethylrhodamine isothiocyanate-dextran, MW = 150,000; T1287, Sigma) was perfused at□9 μl min 1. CSF was sampled at 5 min intervals from cisterna magna as described above for CSF extraction. 60 min into the experiment, the control test solution was replaced with one containing either TRPV4 activator (GSK, 500 nM) or TRPV4 inhibitor (RN1734, 50 μM), with an expected 2-fold ventricular dilution. The fluorescent content was measured in a microplate photometer (545 nm, Synery^TM^ Neo2 Multi-mode Microplate Reader; BioTek Instruments), and the production rate of CSF was calculated from the equation:

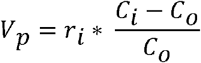

Where *V_p_* = CSF production rate (μl min^-1^), *n* = infusion rate (μl min^-1^), *C_i_* = fluorescence of inflow solution, *C_o_* = fluorescence of outflow solution, calculated based on stable time intervals from 50-65 min and 100-120 min.

### Live imaging of CSF movement

Through a burr hole in the skull of the anaesthetized rat (same coordinates as for ICP and VCP), a Hamilton syringe (RN 0.40, G27, a20, Agntho’s) was placed (4 mm deep) with 15□μl HCO_3_^-^-aCSF containing vehicle (DMSO) or drug of interest (TRPV4 activator, 1 μM GSK; TRPV4 inhibitor, 100 μM RN1734, expected to be immediately diluted in ~ 150 μl native CSF). The content was injected during 10□s to target both lateral ventricles. The procedure was repeated 5 min later, but with addition of the carboxylate dye (MW=1,091, IRDye 800 CW, P/N 929-08972, LI-COR Biosciences) to the syringe content (10μM). The rat was swiftly placed in a Pearl Trilogy Small Animal Imaging System (LI-COR Biosciences) and within 1 min after last ventricular injection, images were obtained at 30 s intervals (800 nm channel, 85 μm resolution, for 5 min). A white field image was taken at the termination of each experiment, after which the rat was sacrificed. The isolated brain was split into the two hemispheres and placed on a coverslip to record a final micrograph to ensure proper targeting of the ventricular compartment (800 nm channel). Images were analysed using LI-COR Image Studio 5.2 (LI-COR Biosciences) and data presented as fluorescence intensity in a region of interest placed in line with lambda, normalized to signals in the first image. Analyses were done in a randomized and blinded fashion.

### RNA-Seq of rat choroid plexus

Isolated rat choroid plexus (lateral and 4^th^) were stored in RNAlater (R0901, Sigma) at −80 °C prior to RNA extraction and library preparation (performed by Novogene Company Limited with NEB Next® Ultra™ RNA Library Prep Kit (NEB)) prior to RNA sequencing (paired-end 150 bp, with 12 Gb output) on an Illumina NovaSeq 6000 (Illumina). Quality control and adapter removal was done with Novogene. The 150 base paired-end reads were mapped to rat reference genome Rnor_6.0.103 *(Rattus norvegicus)* using Spliced Transcripts Alignment to a Reference (STAR) RNA-seq aligner (v.2.7.2a) [63]. The mapped alignment generated by STAR was normalized to transcripts per million (TPM) [64] with RSEM (v 1.3.3) [65]. The RNAseq data obtained from this choroid plexus tissue has also been employed for other analysis [66]. Gene information was gathered with mygene 3.1.0 python library [67, 68] (http://mygene.info) where GO-terms [69–71] for cellular component were extracted. The list of genes annotated as “Voltage-gated ion channels”, “Ligand-gated ion channels”, and “Other Ion channel” was obtained from the Guide to Pharmacology webpage (https://www.guidetopharmacology.org/download.jsp) [72] and employed to generate a list of plasma membrane ion channel proteins (in which ion channels in intracellular membranes were excluded). The list of rat kinases was obtained from the regphos 2.0 database [73], http://140.138.144.141/~RegPhos/). Scripts and program parameters can be found at https://github.com/Sorennorge/MacAulayLab-RNAseq1. Network analysis was carried out with the string database (string-db.org) [74]: Search criteria: Multiple proteins in organism ‘Rattus norvegicus’: TRPV4, SLC12A2 (NKCC1), WNK1, WNK3, WNK4, Stk39 (SPAK).

### Cell surface biotinylation and Western blotting

EZ-link Sulfo-NHS-SS biotin (Thermo Fisher, 1.5 mg/100□μl buffer (125 mM NaCl, 2 mM CaCl_2_, 10 mM triethanolamine, pH 7.5)) was slowly (15 min) injected into the ventricle of an anesthetized rat through a brain infusion cannula (Brain infusion kit 2, Alzet) placed in the rat lateral ventricle as described above for the VCP procedure. The choroid plexus was isolated immediately thereafter and lysed for 30 min in choroid plexus lysis buffer (150 mM NaCl, 5 mM EDTA, 50 mM Tris-HCl, 1% Triton X-100, 0.05% SDS, 0.4 mM pefabloc (Sigma-Aldrich) and 8□μM leupeptin (Sigma-Aldrich)), sonicated at 70% (Sonopuls, Bandelin) and centrifuged at 10,000□×□*g* for 5 min at 4□°C. An aliquot was removed (total fraction) before proceeding with the biotin (luminal) fraction purified on NeutrAvidin (Thermo Fisher) columns (Pierce). The protein concentration was determined with a DC Protein Assay (Bio-Rad) according to the manufacturer’s instruction. 10 μg of total protein was loaded on 4-20% SDS-PAGE gels (Criterion TGX, Bio-rad), before transferred to immobilon-FL membranes (Merck Milipore). Primary and secondary antibodies were diluted 1:1 in Odyssey blocking buffer (LI-COR). The membrane was incubated with the following antibodies: Primary: anti-GAPDH (Millipore #AB2302, 1:5000), and anti-TRPV4 (Alamone #ACC-034, 1:1000). Secondary: 680RD donkey anti-chicken (LI-COR, P/N 925-68075, 1:10.000) and 800CW goat anti-rabbit (LI-COR, P/N 926-32211, 1:10.000). The final blots were visualized by the Odyssey CLx imaging system and analyzed with Image Studio 5.2.5 (LI-COR).

### Tissue preparation and immunohistochemistry

Anesthetized rats were perfusion fixed with 4% paraformaldehyde and the brains removed for post-fixation in the same fixative overnight, cryoprotected in 25% sucrose, and frozen on solid CO_2_. Saggital and coronal sections (16 μm) were cut and stored at −20°C. Blockage was done with 10% swine serum diluted in PBS+1%Tween-20 (PBST), and the sections were immunolabeled with primary rabbit anti-NKCC1 (Abcam #AB59791, 1:400) and mouse anti-TRPV4 (BD BioSciences #AB53079, 1:500) overnight at 4°C, and Alexa Fluor™488 and Alexa Fluor™647 (ThermoFisher Scientific 1:500) for 2 h at room temperature. Finally, the sections were mounted with ProLong Gold DAPI mounting medium (Dako). The confocal images were acquired with a Zeiss LSM710 point laser (Argon Lasos RMC781272) scanning confocal microscope with a Zeiss Plan-Apochromat 63 ×/numerical aperture (NA) 1.4 oil objective (Carl Zeiss, Oberkochen). All micrographs were sampled in a frame scan mode.

### Proximity Ligation Assay

Proximity ligation assay by indirect detection was done using Duolink reagents and instructions (Sigma-Aldrich) on rat brain cryosections (16 μm) (see ‘Tissue preparation’ section). After incubation in Duolink blocking solution, the tissue sections were incubated with anti-TRPV4 (LS-Bio #LS-C94498, 1:400) and anti-NKCC1 (Dundee University #S022D, 2μg/ml) overnight at 4°C. The secondary antibodies were conjugated to a MINUS and a PLUS PLA probe, and were detectable as fluorescent speckles only upon close contact [40]. A technical control omitting both of the primary antibodies was included. Ligation and amplification of the samples were done according to the Duolink protocol, the sections were mounted with coverslips using the Duolink In Situ Mounting Medium with DAPI (Sigma-Aldrich). Micrographs were acquired with a Zeiss LSM700 point laser (Argon Lasos RMC781272) scanning confocal microscope with a Zeiss Plan-Apochromat 20×/numerical aperture (NA) 1.6 oil objective (Carl Zeiss, Oberkochen). All micrographs were sampled in a frame scan mode.

### Calcium imaging in isolated choroid plexuses

Acutely isolated choroid plexuses were mounted on glass coverslips coated with Cell-Tak® (Corning; 1 part 2 M Na2CO3:9 parts Cell-Tak® with 2% isopropanol). The coverslip was placed in an open POCmini2 perfusion chamber (PeCon) in a heating chamber (37°C) in which the choroid plexus was loaded with Fluo-8 AM (5 μM, 30 min, Abcam, ab142773) with subsequent continuous wash with HEPES-aCSF (~1 ml/min) prior to inclusion of TRPV4 modulators (200 μM RN or 1 μM GSK). Live imaging was performed with an inverted confocal microscope (Carl Zeiss, Cell-Observer, LED, 10x 0.3 NA EC Plan Neofluar objective, ZEN black ed. software) and the analysis carried out in a randomized and blinded fashion.

### ^86^Rb^+^ efflux experiments

Acutely isolated lateral choroid plexuses were placed in cold HCO_3_^-^-aCSF but allowed to recover at 37°C for 5–10□min before the experiment. Choroidal isotope accumulation was performed by a 10 min incubation in gas-equilibrated HCO_3_^-^-aCSF with 1μCi/ml ^86^Rb^+^ (NEZ07200 and 4□μCi/ml ^3^H-mannitol (NET101, extracellular marker), from PerkinElmer and Polatom). The choroid plexus was briefly washed (15□s) prior to incubation in 0.5 ml efflux medium (HCO_3_^-^-aCSF containing the indicated modulators; 2MμM bumetanide (NKCC1 inhibitor), 100 nM GSK (TRPV4 agonist), 50 μM RN (TRPV4 inhibitor), 20 μM Closantel (SPAK kinase-inhibitor), 20 nM WNK463 (WNK kinase-inhibitor), 25 μM LPA, or vehicle (DMSO). 0.2ml of the efflux medium was collected into scintillation vials at 10□s time intervals and replaced with fresh HCO_3_^-^-aCSF. Upon termination of the experiment, the choroid plexuses were dissolved in 1□ml Solvable (6NE9100, PerkinElmer) and the isotope content determined by liquid scintillation counting with Ultima Gold™ XR scintillation liquid (60133119. PerkinElmer) in a Tri-Carb 2900TR Liquid Scintillation Analyzer (Packard). The choroid plexus ^86^Rb^+^ content corrected for ^3^H-mannitol content (extracellular background) was calculated for each time point, and the natural logarithm of the choroid plexus content *A_t_*/*A*_0_ was plotted against time. Slopes indicating the ^86^Rb^+^ efflux rate constants (min^-1^) were determined from linear regression analysis [25, 75].

### Electrophysiological recordings in heterologously expressing *Xenopus laevis* oocytes

Defolliculated oocytes obtained from *Xenopus laevis* were purchased from Ecocyte Bioscience. cRNA (TRPV4) was prepared from linearized plasmids (pXOOM containing cDNA encoding TRPV4 using the mMESSAGE mMACHINE T7 kit (Ambion) and extracted with MEGAclear (Ambion), according to the manufacturer’s instructions. The oocytes were microinjected with 4□ng cRNA per oocyte with a Nanoject microinjector (Drummond Scientific Company). The oocytes were kept at 19°C for 3 days in Kulori medium prior to experiments. Oocytes were kept in Kulori medium (90 mM NaCl, 1 mM KCl, 1 mM CaCl_2_, 1 mM MgCl_2_, 5 mM HEPES (pH 7.4)) with inclusion of ruthenium red (100 μm; Sigma Aldrich, R-2751) to suppress TRPV4 activity and ensuing cell death [76]. Conventional two-electrode voltage-clamp was performed using a DAGAN CA-1B High Performance oocyte clamp (DAGAN, Minneapolis, MN, USA) with a Digidata 1440A interface controlled by pCLAMP software, version 10.5 (Molecular Devices, Burlingame, CA, USA). Borosilicate glass capillaries were used to pull electrodes (PIP5; HEKA Elektronik, Lambrecht, Germany) with a resistance of 1.5–3 MΩ when filled with 1 m KCl. Oocytes were placed in an experimental recording chamber and perfused with a test solution containing 100 mM NaCl, 2 mM KCl, 1 mM MgCl_2_, 1 mM CaCl_2_ and 10 mM HEPES (Tris buffered pH 7.4, 213 mosmol 1^-1^). Current traces were obtained by stepping the holding potential of −20 mV to test potentials ranging from −130 mV to +50 mV in increments of 15 mV in pulses of 200 msec. Recordings were low pass-filtered at 500 Hz, sampled at 1 kHz and the steadystate current activity was analysed at 160–180 ms after applying the test pulse.

## Data Availability

We confirm that all data from this study will be available upon request.

## REFERENCES

1. Spector, R., et al., A balanced view of choroid plexus structure and function: Focus on adult humans. Exp Neurol, 2015. 267: p. 78–86.

2. MacAulay, N., Molecular mechanisms of brain water transport. Nat Rev Neurosci, 2021. 22(6): p. 326–344.

3. Johanson, C.E., et al., Multiplicity of cerebrospinal fluid functions: New challenges in health and disease. Cerebrospinal Fluid Res, 2008. 5: p. 10.

4. Serot, J.M., J. Zmudka, and P. Jouanny, A possible role for CSF turnover and choroid plexus in the pathogenesis of late onset Alzheimer’s disease. J Alzheimers Dis, 2012. 30(1): p. 17–26.

5. Hallaert, G.G., et al., Endoscopic coagulation of choroid plexus hyperplasia. J Neurosurg Pediatr, 2012. 9(2): p. 169–77.

6. Karimy, J.K., et al., Inflammation-dependent cerebrospinal fluid hypersecretion by the choroid plexus epithelium in posthemorrhagic hydrocephalus. Nat Med, 2017. 23(8): p. 997–1003.

7. Ducros, A. and V. Biousse, Headache arising from idiopathic changes in CSF pressure. Lancet Neurol, 2015. 14(6): p. 655–68.

8. Chen, Q., et al., Post-hemorrhagic hydrocephalus: Recent advances and new therapeutic insights. J Neurol Sci, 2017. 375: p. 220–230.

9. Strahle, J., et al., Mechanisms of hydrocephalus after neonatal and adult intraventricular hemorrhage. Transl Stroke Res, 2012. 3(SuppI 1): p. 25–38.

10. Persson, E.K., et al., Hydrocephalus in children born in 1999-2002: epidemiology, outcome and ophthalmological findings. Childs Nerv Syst, 2007. 23(10): p. 1111–8.

11. Kahle, K.T., et al., Hydrocephalus in children. Lancet, 2016. 387(10020): p. 788–99.

12. Abdelmalik, P.A. and W.C. Ziai, Spontaneous Intraventricular Hemorrhage: When Should Intraventricular tPA Be Considered? Semin Respir Crit Care Med, 2017. 38(6): p. 745–759.

13. Roeder, S.S., et al., Influence of the Extent of Intraventricular Hemorrhage on Functional Outcome and Mortality in Intracerebral Hemorrhage. Cerebrovasc Dis, 2019. 47(5-6): p. 245–252.

14. Garton, T., et al., Challenges for intraventricular hemorrhage research and emerging therapeutic targets. Expert Opin Ther Targets, 2017. 21(12): p. 1111–1122.

15. Starnoni, D., et al., Thrombolysis for non-traumatic intra-ventricular hemorrhage in adults: a critical reappraisal. Minerva Anestesiol, 2017. 83(9): p. 982–993.

16. Iwaasa, M., et al., Safety and feasibility of combined coiling and neuroendoscopy for better outcomes in the treatment of severe subarachnoid hemorrhage accompanied by massive intraventricular hemorrhage. J Clin Neurosci, 2013. 20(9): p. 1264–8.

17. Hokari, M., et al., Ruptured aneurysms of the choroidal branches of the posterior inferior cerebellar artery: a review of the literature and a case report. Acta Neurochir (Wien), 2010. 152(3): p. 515–8.

18. McLaughlin, N. and M.W. Bojanowski, Ruptured aneurysm at the choroidal branch of the posterior inferior cerebellar artery: a case report, review of the literature and proposed pathogenesis. Br J Neurosurg, 2005. 19(3): p. 250–3.

19. Naff, N.J. and S. Tuhrim, Intraventricular hemorrhage in adults: complications and treatment. New Horiz, 1997. 5(4): p. 359–63.

20. Klebe, D., et al., Posthemorrhagic hydrocephalus development after germinal matrix hemorrhage: Established mechanisms and proposed pathways. J Neurosci Res, 2020. 98(1): p. 105–120.

21. Hill, A., G.D. Shackelford, and J.J. Volpe, A potential mechanism of pathogenesis for early posthemorrhagic hydrocephalus in the premature newborn. Pediatrics, 1984. 73(1): p. 19–21.

22. Mazzola, C.A., et al., Pediatric hydrocephalus: systematic literature review and evidencebased guidelines. Part 2: Management of posthemorrhagic hydrocephalus in premature infants. J Neurosurg Pediatr, 2014. 14 Suppl 1: p. 8–23.

23. Shooman, D., H. Portess, and O. Sparrow, A review of the current treatment methods for posthaemorrhagic hydrocephalus of infants. Cerebrospinal Fluid Res, 2009. 6: p. 1.

24. Eisenberg, H.M., J.G. McComb, and A.V. Lorenzo, Cerebrospinal fluid overproduction and hydrocephalus associated with choroid plexus papilloma. J Neurosurg, 1974. 40(3): p. 381–5.

25. Steffensen, A.B., et al., Cotransporter-mediated water transport underlying cerebrospinal fluid formation. Nat Commun, 2018. 9(1): p. 2167.

26. Javaheri, S. and K.R. Wagner, Bumetanide decreases canine cerebrospinal fluid production. In vivo evidence for Nad cotransport in the central nervous system. J Clin Invest, 1993. 92(5): p. 2257–61.

27. Oernbo, E.K., et al., Cerebrospinal fluid formation is controlled by membrane transporters to modulate intracranial pressure. bioRxiv, 2021: p. 2021.12.10.472067.

28. Lummis, N.C., et al., LPA1/3 overactivation induces neonatal posthemorrhagic hydrocephalus through ependymal loss and ciliary dysfunction. Sci Adv, 2019. 5(10): p. eaax2011.

29. Yung, Y.C., et al., Lysophosphatidic acid signaling may initiate fetal hydrocephalus. Sci Transl Med, 2011. 3(99): p. 99ra87.

30. Crack, P.J., et al., Anti-lysophosphatidic acid antibodies improve traumatic brain injury outcomes. J Neuroinflammation, 2014. 11: p. 37.

31. Aoki, J., et al., Serum lysophosphatidic acid is produced through diverse phospholipase pathways. J Biol Chem, 2002. 277(50): p. 48737–44.

32. Liedtke, W., et al., Vanilloid receptor-related osmotically activated channel (VR-OAC), a candidate vertebrate osmoreceptor. Cell, 2000. 103(3): p. 525–35.

33. Takayama, Y., et al., Modulation of water efflux through functional interaction between TRPV4 and TMEM16A/anoctamin 1. FASEB J, 2014. 28(5): p. 2238–48.

34. Toft-Bertelsen, T.L. and N. MacAulay, TRPing to the Point of Clarity: Understanding the Function of the Complex TRPV4 Ion Channel. Cells, 2021. 10(1).

35. Preston, D., et al., Activation of TRPV4 stimulates transepithelial ion flux in a porcine choroid plexus cell line. Am J Physiol Cell Physiol, 2018. 315(3): p. C357–C366.

36. Simpson, S., et al., Cytokine and inflammatory mediator effects on TRPV4 function in choroid plexus epithelial cells. Am J Physiol Cell Physiol, 2019. 317(5): p. C88l–C893.

37. Hochstetler, A.E., et al., TRPV4 antagonists ameliorate ventriculomegaly in a rat model of hydrocephalus. JCI Insight, 2020. 5(18).

38. Beaudet, L., et al., AlphaLISA immunoassays: the no-wash alternative to ELISAs for research and drug discovery. Nature Methods, 2008. 5(12): p. an8–an9.

39. Vincent, F., et al., Identification and characterization of novel TRPV4 modulators. Biochem Biophys Res Commun, 2009. 389(3): p. 490–4.

40. Soderberg, O., et al., Direct observation of individual endogenous protein complexes in situ by proximity ligation. Nat Methods, 2006. 3(12): p. 995–1000.

41. Lykke, K., et al., The search for NKCC1-selective drugs for the treatment of epilepsy: Structure-function relationship of bumetanide and various bumetanide derivatives in inhibiting the human cation-chloride cotransporter NKCC1A. Epilepsy Behav, 2016. 59: p. 42–9.

42. Park, S., et al., Ca(2+) is a Regulator of the WNK/OSR1/NKCC Pathway in a Human Salivary Gland Cell Line. Korean J Physiol Pharmacol, 2015. 19(3): p. 249–55.

43. Zhang, J., X. Deng, and K.T. Kahle, Leveraging unigue structural characteristics of WNK kinases to achieve therapeutic inhibition. Sci Signal, 2016. 9(450): p. e3.

44. Cinalli, G., et al., Hydrocephalus in agueductal stenosis. Childs Nerv Syst, 2011. 27(10): p. 1621–42.

45. Beni-Adani, L, et al., The occurrence of obstructive vs absorptive hydrocephalus in newborns and infants: relevance to treatment choices. Childs Nerv Syst, 2006. 22(12): p. 1543–63.

46. Oi, S. and C. Di Rocco, Proposal of “evolution theory in cerebrospinal fluid dynamics” and minor pathway hydrocephalus in developing immature brain. Childs Nerv Syst, 2006. 22(7): p. 662–9.

47. Rekate, H.L., The definition and classification of hydrocephalus: a personal recommendation to stimulate debate. Cerebrospinal Fluid Res, 2008. 5: p. 2.

48. Karimy, J.K., et al., Inflammation in acquired hydrocephalus: pathogenic mechanisms and therapeutic targets. Nat Rev Neurol, 2020. 16(5): p. 285–296.

49. Relkin, N., et al., Diagnosing idiopathic normal-pressure hydrocephalus. Neurosurgery, 2005. 57(3 Suppl): p. S4–16; discussion ii-v.

50. Williams, M.A. and J. Malm, Diagnosis and Treatment of Idiopathic Normal Pressure Hydrocephalus. Continuum (Minneap Minn), 2016. 22(2 Dementia): p. 579–99.

51. Lolansen, S.D., et al., Inflammatory Markers in Cerebrospinal Fluid from Patients with Hydrocephalus: A Systematic Literature Review. Dis Markers, 2021. 2021: p. 8834822.

52. Yung, Y.C., N.C. Stoddard, and J. Chun, LPA receptor signaling: pharmacology, physiology, and pathophysiology. J Lipid Res, 2014. 55(7): p. 1192–214.

53. Atlas, A.B., http://www.brain-map.org.

54. Bothwell, S.W., et al., CSF Secretion Is Not Altered by NKCC1 Nor TRPV4 Antagonism in Healthy Rats. Brain Sci, 2021. 11(9).

55. Kahle, K.T., J. Rinehart, and R.P. Lifton, Phosphoregulation of the Na-K-2CI and K-CI cotransporters by the WNK kinases. Biochim Biophys Acta, 2010. 1802(12): p. 1150–8.

56. Piechotta, K., J. Lu, and E. Delpire, Cation chloride cotransporters interact with the stress-related kinases Ste20-related proline-alanine-rich kinase (SPAK) and oxidative stress response 1 (OSR1). J Biol Chem, 2002. 277(52): p. 50812–9.

57. Zhang, J., et al., Modulation of brain cation-CI(-) cotransport via the SPAK kinase inhibitor ZT-1a. Nat Commun, 2020. 11(1): p. 78.

58. Huang, H., et al., The WNK-SPAK/OSR1 Kinases and the Cation-Chloride Cotransporters as Therapeutic Targets for Neurological Diseases. Aging Dis, 2019. 10(3): p. 626–636.

59. Tanioka, H., et al., Three-dimensional reconstructed MR imaging of the inner ear. Radiology, 1991. 178(1): p. 141–4.

60. Avants, B.B., et al., The Insight ToolKit image registration framework. Front Neuroinform, 2014. 8: p.44.

61. Tustison, N.J., et al., N4ITK: improved N3 bias correction. IEEE Trans Med Imaging, 2010. 29(6): p. 1310–20.

62. Yushkevich, P.A., et al., User-guided 3D active contour segmentation of anatomical structures: significantly improved efficiency and reliability. Neuroimage, 2006. 31(3): p. 1116–28.

63. Dobin, A., et al., STAR: ultrafast universal RNA-seq aligner. Bioinformatics, 2013. 29(1): p. 15–21.

64. Abrams, Z.B., et al., A protocol to evaluate RNA sequencing normalization methods. BMC Bioinformatics, 2019. 2O(SuppI 24): p. 679.

65. Li, B. and C.N. Dewey, RSEM: accurate transcript quantification from RNA-Seq data with or without a reference genome. BMC Bioinformatics, 2011. 12: p. 323.

66. Lolansen, S.D., et al., Elevated CSF inflammatory markers in patients with idiopathic normal pressure hydrocephalus do not promote NKCC1 hyperactivity in rat choroid plexus. Fluids Barriers CNS, 2021. 18(1): p. 54.

67. Xin, J., et al., High-performance web services for querying gene and variant annotation. Genome Biol, 2016. 17(1): p. 91.

68. Wu, C., I. Macleod, and A.I. Su, BioGPS and MyGene.info: organizing online, gene-centric information. Nucleic Acids Res, 2013. 4l(Database issue): p. D561–5.

69. Carbon, S., et al., AmiGO: online access to ontology and annotation data. Bioinformatics, 2009. 25(2): p. 288–9.

70. Day-Richter, J., et al., OBO-Edit--an ontology editor for biologists. Bioinformatics, 2007. 23(16): p. 2198–200.

71. Mi, H., et al., PANTHER version 14: more genomes, a new PANTHER GO-slim and improvements in enrichment analysis tools. Nucleic Acids Res, 2019. 47(D1): p. D419–D426.

72. Alexander, S.P., et al., THE CONCISE GUIDE TO PHARMACOLOGY 2021/22: Ion channels. Br J Pharmacol, 2021. 178 Suppl 1: p. S157–S245.

73. Huang, K.Y., et al., RegPhos 2.0: an updated resource to explore protein kinase-substrate phosphorylation networks in mammals. Database (Oxford), 2014. 2014(0): p. bau034.

74. Szklarczyk, D., et al., STRING v11: protein-protein association networks with increased coverage, supporting functional discovery in genome-wide experimental datasets. Nucleic Acids Res, 2019. 47(D1): p. D607–D613.

75. Keep, R.F., J. Xiang, and A.L. Betz, Potassium cotransport at the rat choroid plexus. Am J Physiol, 1994. 267(6 Pt 1): p. C1616–22.

76. Toft-Bertelsen, T.L., D. Krizaj, and N. MacAulay, When size matters: transient receptor potential vanilloid 4 channel as a volume-sensor rather than an osmosensor. J Physiol, 2017. 595(11): p. 3287–3302.

